# Modular, reconfigurable fiber-based neural probe (MoRF probe) with interchangeable and tunable optical waveguide, microfluidic channel, and microelectrodes

**DOI:** 10.1101/2025.06.29.662251

**Authors:** Hengji Huang, Yue Liu, Shuo Yang, Song Hu, Daniel Fine English, Xiaoting Jia

**Author notes:** These authors contributed equally to this work (Hengji Huang, Yue Liu).

## Abstract

Neuroscience is at an exciting juncture at which large-scale *in vivo* electrophysiology, including in humans, is intersecting symbiotically with revolutionary computational and statistical methods. Maintaining this relationship requires increasingly advanced neural probes for multimodal interrogation of neural circuitry (e.g. perturbation experiments are of value to computational modeling). Here, we developed a low cost modular and reconfigurable recording and stimulation fiber-based neural probe (MoRF) fabricated via a first step thermal drawing process (TDP) and a second step thermal tapering process (TTP). We demonstrated the device modularity and reconfigurability in several functional variations of the same device, as well as the ability to adjust the distance between different sensing elements. We validated the electrical, optical and microfluidic drug delivery performance of the MoRF probe and demonstrated its *in vivo* electrophysiological recording and optogenetic stimulation capabilities in awake mice.

## Introduction

Bi-directional neural interfaces are essential tools for uncovering the complexities of the brain. By manipulating neuronal function and recording neural activity, neural interfaces provide researchers with the means to interrogate neural circuit mechanisms, find treatment for neurological disorders, and develop brain machine interfaces. A diverse set of impressive tools for recording and modulating neural activity have recently been developed and adopted across many fields of inquiry. These include electrophysiological recording that enables high temporal resolution monitoring of neuronal populations^1,2^, Optogenetic excitation that allows for cell-type specific neural modulation with high spatial resolution^3,4^, and localized pharmacological modulation (e.g. to probe the function of neuromodulator receptors in neural circuits)^5^. A device that multiplexes these modalities would be a powerful platform for multifaceted interrogation of neural circuitry useful for addressing core questions such as causal relationships between neural circuit activity and physical behavior of organisms.

Although technological advancements have resulted in the creation of increasingly sophisticated and effective multimodal probes, customization, modularity, and reconfigurability still pose significant challenges. Current state-of-the-art multimodal devices have impressive abilities but their functionalities and geometrical arrangement cannot be altered post-fabrication. For example, Yan et al. introduced the origami probe whose sensing element arrays with different functionalities were fabricated individually and stacked together to form a multifunctional device^19^. This innovative method significantly enhances modularity at the fabrication stage. However, the device’s functions are fixed after the probe has been fabricated. Ideally a probe would be customizable and reconfigurable not just post-fabrication but also post-implantation into the brain. Such a feature is essential for neuroscientific studies that require different device modalities at different phases of the experiment. It can also prove extremely useful in chronic implantations where device failure, glial scarring and degradation of device performance from micromovements causing electrodes to lose units over time undermines recording quality^20–25^. Various partial solutions to enhance chronic recording quality have been proposed, such as the addition of biocompatible coatings to mitigate device degradation and microdrives to reposition the probe when units are lost^26,27^. A post-implantation reconfigurable device offers a novel alternative solution. Without advancing or retracting the entire probe, as would be the case with wire or Si probes, the center electrode set would independently move inside the whole probe body; the probe body remaining stable is important for overall device stability. Such a modular, reconfigurable neural interface that can be customized at will to specific experimental requirements without dependence on cleanroom fabrication offers a cost-effective and adaptable platform for advancing research ranging from basic neuroscience to clinical neuroengineering.

In this work, we address this gap by developing a Modular, Reconfigurable Fiber-based probe (MoRF probe) using a thermal drawing process (TDP) and a thermal tapering process (TTP) as first and second fabrication steps. TDP is a promising scalable method for fabricating flexible multifunctional neural probes^16,28–32^ that starts by fabricating a macro scale preform (a scaled-up version of the device). It is then heated and thermally drawn into hundreds of meters of thin (100-400 µm) fibers, which are cut into short sections and connectorized to form neural probes. This process enables highly scalable fabrication by producing thousands of devices from one preform without needing a cleanroom, which significantly minimizes fabrication time and cost compared to other common fabrication methods. In addition, neural probes require reliable low-impedance connectorization between the implanted device and the back-end interface. The necessarily small size of probes makes it a significant challenge to connect high-density functional features in the fibers with various geometrical arrangements to external recording and stimulation instruments. To solve this problem, we developed a new thermal tapering process (TTP) to complement TDP. Here instead of the standard process of using TDP to draw the probe in a single step, we use TDP to first draw an intermediate-size structure called a mini-preform. TTP then heats and pulls the mini-preform to form a tapered structure with a large backend for easy connection, and a small sensing end for minimal damage during insertion.

Drawing inspiration from the 200 million year old mosquito proboscis, which is a tube-within-a-tube comprised of a bundle of movable stylets with different functions, the MoRF probe consists of a cylindrical “outer” probe with a hole in the center through which an “inner” probe can be inserted and then adjusted or fixed in place. Such a design allows the user to create custom and dynamic probes with the ability for users to assemble the necessary complement of elements needed for a specific experiment. Here we show the versatility of our probe by demonstrating several possible configurations with different functionalities, including optogenetic stimulation and multi-site drug delivery and enhanced multi-site 32 channel electrical recording. We also demonstrate our probe’s compatibility with sidewall electrode exposure via femtosecond laser etching, which can be used to further enhance its electrophysiological recording capabilities^30^. Our probe’s electrophysiological recording, optogenetic stimulation, and multi-site, multi-drug delivery functionalities are validated via impedance measurements and implantation simulations in 0.6% agar tissue phantom. Electrical recording and optical stimulation capabilities are then tested via *in vivo* implantation and recording in awake and behaving mice.

## Results

### Preform Fabrication and Tapering

The device is fabricated via a combination of Thermal Drawing Process (TDP) and Thermal Tapering Process (TTP) as demonstrated in Fig.1a-c. The preform was fabricated via a series of thermal consolidation and machining steps. For the outer probe’s preform, polycarbonate (PC) sheets were wrapped around a Teflon rod and consolidated in a vacuum furnace. After that, the Teflon rod was removed to form a hollow PC tube. A Computer Numerical Control (CNC) machine was then used to machine 16 equally spaced hollow channels into the tube. The hollow channels were then filled with BiSn alloy strips and covered with PC film. The preform was then consolidated again under vacuum. Next, the preform was mounted in a 3-zone (top, middle, bottom) gradient furnace, heated until softened and thermally drawn into 2 mm diameter mini-preforms. Preform and mini-preform fabrication for different inner probe configurations were also similar should the inner probe not be commercially available.

**Fig. 1:**
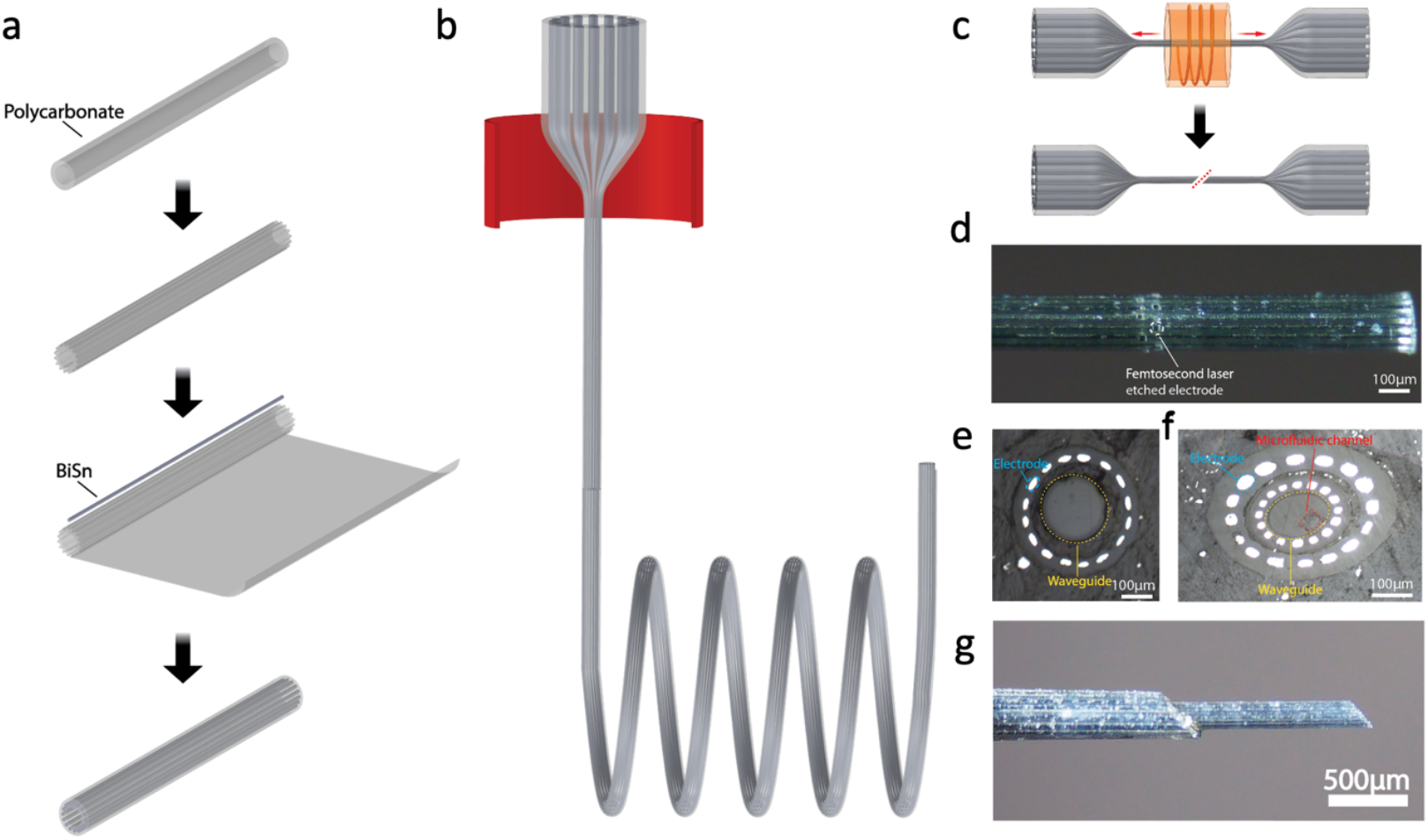
MoRF probe fabricated using TDP and TTP. **a**. Schematic of the preform fabrication process. **b**. Illustration of the TDP in which 30 mm diameter preform is heated and pulled into 2 mm diameter mini-preform. **c**. Illustration of the TTP of mini-preform. Mini-preform is mounted on a TTP setup, heated until softened, tapered, and cut in the middle at a desired angle, resulting in two individual probes. **d**. Side-view optical image of a 16-channel probe with side wall electrodes exposed by femtosecond laser micromachining. **e**. Cross-sectional image of a 16-channel configuration which consists of sixteen electrodes and one silica optical waveguide. **f**. Cross-sectional images of a 32-channel configuration which consists of thirty-two electrodes, one microfluidic channel, and one polymer optical waveguide. **g**. Side-view optical image of the 32-channel probe with a fiber-in-fiber configuration: an inner fiber with 16-electrodes, 1 waveguide, and 1 microfluidic channel is inserted inside the hollow core of a 16-electrode outer fiber.

The mini-preform was then loaded into a custom-built TTP setup, tapered, and cut in the middle to form two devices. The outer probe’s dimensions were minimized while still maintaining a large enough hollow core for the insertion of the inner probe. The inner probe’s length was adjusted to ensure partial extension of the inner probe’s tip beyond the outer probe for the fully assembled device. An image of a device with electrodes exposed along the probe sidewall via femtosecond laser micromachining is shown in Fig.1d. Polished cross sections of different device configurations (16 electrodes and 1 movable silica waveguide; 32 electrodes with the inner 16 electrodes movable, 1 movable polymer waveguide, and 1 drug delivery channel) are shown in Fig.1e and Fig.1f, respectively. Fig. 1g shows the side-view optical image of a 32-channel probe with a fiber-in-fiber configuration. An inner fiber with 16 electrodes, 1 waveguide, and 1 microfluidic channel is inserted inside the hollow core of an outer fiber. The total diameter of the device is ~300 µm, with the tip (inner fiber) diameter of ~150 µm.

### Probe Connectorization & Assembly

Connecting the probe’s sensing elements was accomplished on the large backend of the tapered devices similar to how it was done in our previous publication^5^, as is shown in Fig.2a. The probe was electrically connected by heating the probe’s backend to the melting point of BiSn (150°C) and inserting copper wires in the BiSn electrodes. In configurations with silica waveguides, standard 1.25 mm optical fiber ferrules (Thorlabs, Inc.) were mounted onto the waveguides. For configurations with drug delivery channels, the microfluidic connections were accomplished by inserting a thermally drawn thin PC tubing into the microfluidic channel and sealing the connection with UV resin. The outer probe was then attached to a custom designed PCB via UV resin and the electrical connections soldered on. The inner probe was inserted into the central hollow channel and attached to the movable section of a metal microdrive (RD2drive, 3Dneuro) via either UV resin or polyethylene glycol (PEG) depending on whether probe reconfigurability is required. An example of a fully connected probe is shown in Fig.2b. To maximize the fully assembled probe’s sensing element reconfigurability, the microdrive was adjusted to provide ample room for the inner probe to both extend out from and retract into the outer probe. The immovable base of the microdrive was then attached to the outer probe by UV resin. The microdrive can then be leveraged to adjust the height difference between the inner and outer probes. Examples of the final device’s sensing element reconfiguration capabilities are demonstrated in Fig.2c where the inner probe (a silica waveguide) height was adjusted via the microdrive. It is to be noted that the microdrive can also be opted out, and the inner fiber can be attached to the outer fiber by inserting a drug delivery tube into the gap and sealing it with an adhesive. This can provide an extra drug delivery channel by utilizing the gap between the inner and outer probe at the cost of probe reconfigurability.

**Fig. 2:**
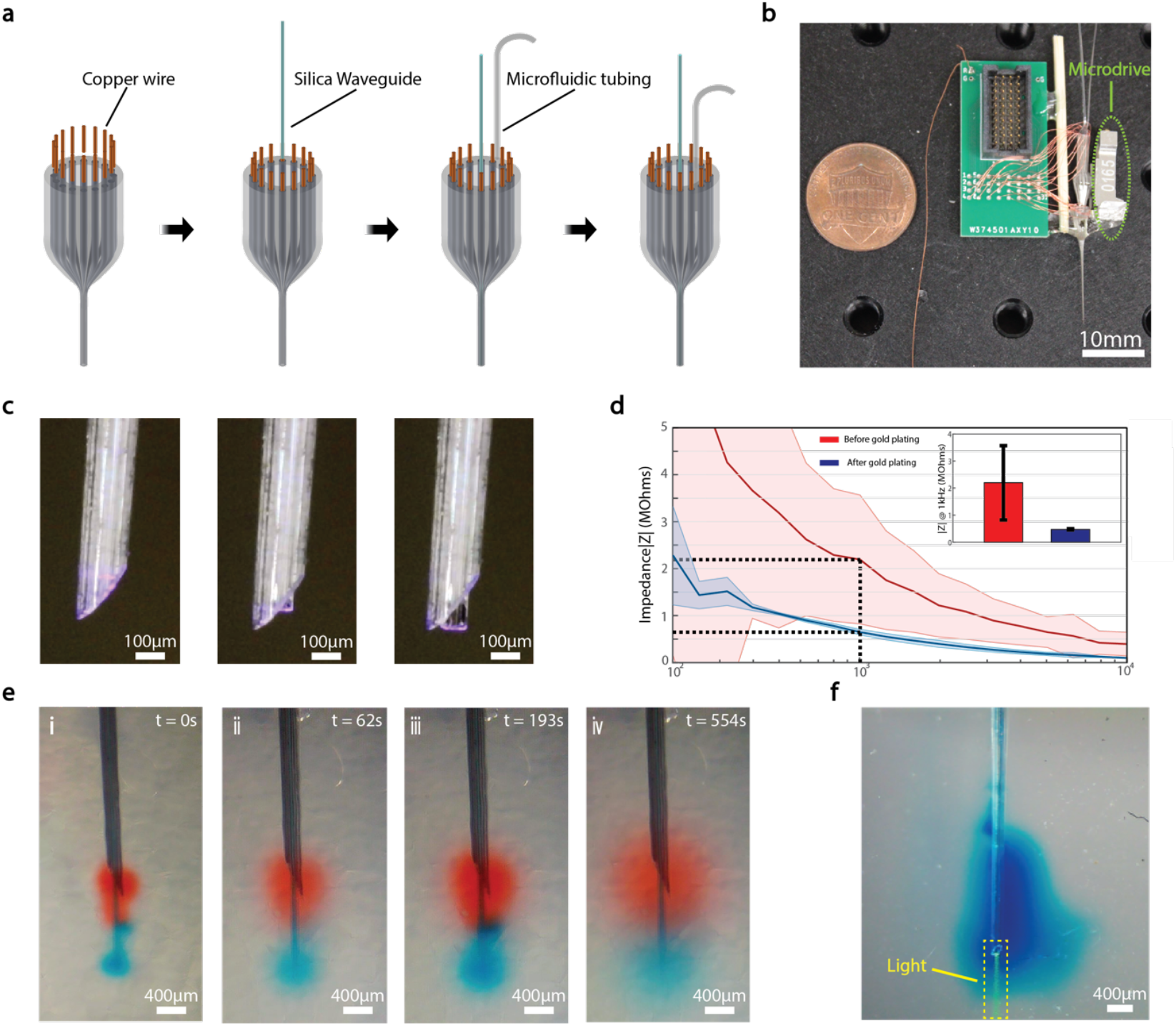
MoRF probe connectorization and characterization. **a**. Schematic of the connectorization process. **b**. Photograph of a fully connected probe with thirty-two electrodes, one waveguide and one drug delivery channel. **c**. Photograph demonstrating probe sensing element reconfiguration. **d**. Impedance of BiSn electrodes before (red) and after (blue) gold plating. **e**. Time lapse photos demonstrating multisite multi-drug injection in 0.6% agarose gel (red, blue food dye coloring) **f**. Photograph demonstrating the probe’s multisite optical stimulation and drug delivery capability in an agarose gel.

### Polishing and Gold Plating

The device was then polished at a 45° angle via an optical fiber polisher (Krell tech) to achieve additional depth distribution for the probe’s sensing elements and minimize tissue damage during probe implantation. To optimize our device’s electrophysiology performance, gold was electroplated on the electrodes by cyclic chronoamperometry using a potentiostat (Gamry Instruments Inc.) and gold Non-Cyanide solution.

### Probe characterization

The device’s electrical characterization was performed via electrochemical impedance spectroscopy (EIS) in saline using a potentiostat. Gold plating consistently lowered the impedance of the electrodes to below 1 MΩ, a practical cutoff for reliably detecting single unit activities *in vivo*. Electrode impedances (n=4) in saline before and after gold plating are shown in Fig.2d. Our device’s multisite multi-drug delivery capabilities were demonstrated by simultaneously injecting two different food dyes at different heights in agarose gel as shown in Fig.2e. Similarly, optical delivery capabilities were demonstrated in 0.6% agarose gel as shown in Fig.2f.

### In vivo electrophysiology recording

Electrophysiological recording capabilities were demonstrated via acute recording experiments from head-fixed awake mice voluntarily navigating a 1D virtual reality environment on a treadmill b (n=3). A 16-channel device was slowly inserted via a linear translation stage into the brain, targeting the dorsal CA1 region of the hippocampus. In local field potential recordings within ~100 µm of the pyramidal layer, we observed normal physiological activity including characteristic oscillations such as the theta oscillation (4-12 Hz) and sharp wave-ripples (100-250Hz). Fig.3a-c shows example wideband (0.1-6000Hz) extracellular traces from a recording session, demonstrating persistent theta oscillations during ambulation and sharp wave ripples (SPW-Rs) in rest. Multiunit activity was also clear in these same channels. In Fig. 3d, the power spectrogram shows neural activity in the frequency domain (frequency bins, 0.1-300Hz) throughout the recording session. We also identified four putative single units via spike sorting from one recording session. Fig.3e shows for each unit the spike train autocorrelation histogram, cross-correlation histogram and the mean waveform. These four units can putatively be classified based on the shape of the waveform and spike train autocorrelation features (putative ID, blue: pyramidal cell, orange: interneuron, green: pyramidal cell, and purple: interneuron). These results demonstrate our device’s ability to acquire both single-unit and local field potential (LFP) activities without disrupting local circuitry in awake behaving mice.

**Fig. 3:**
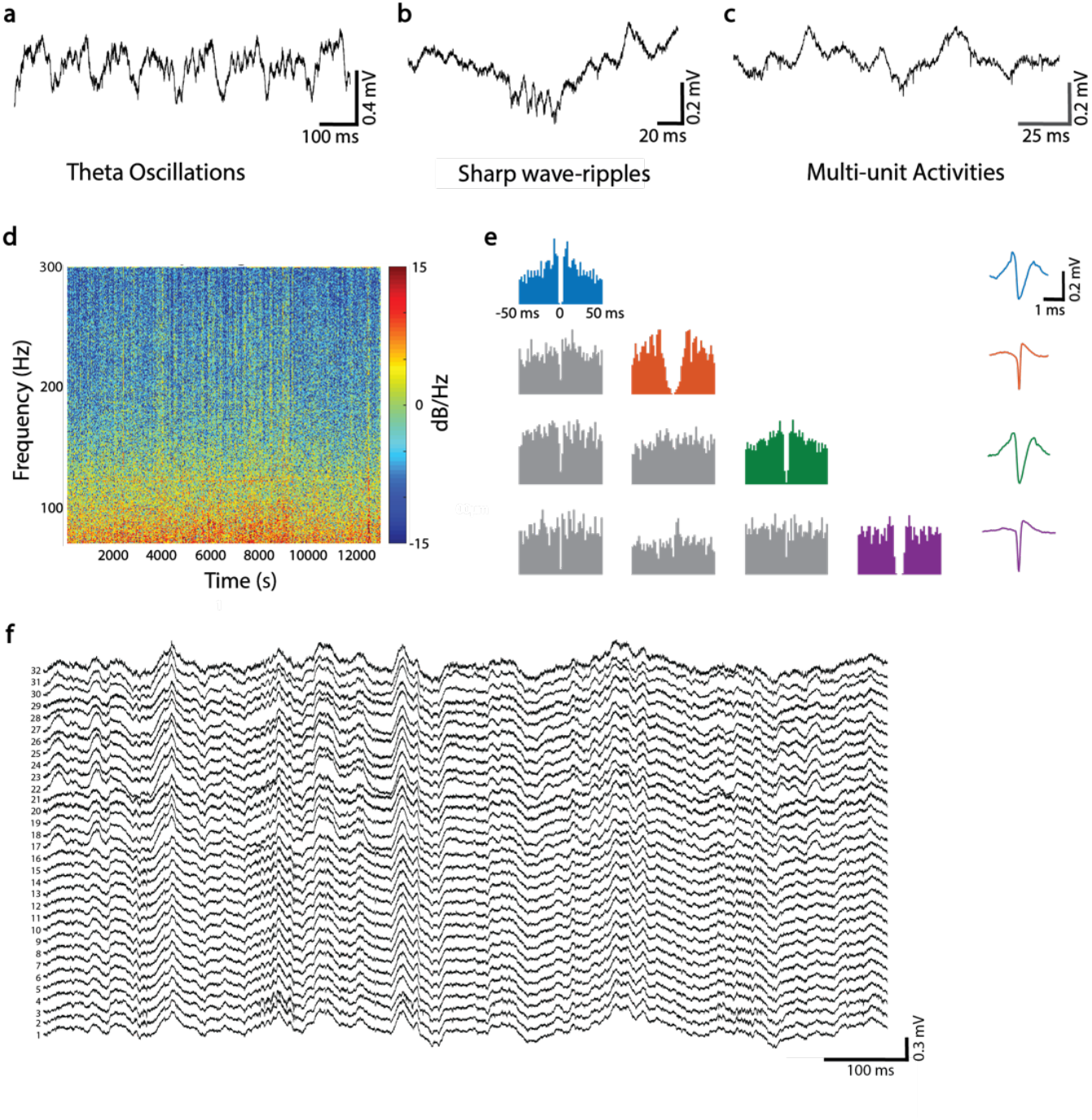
In vivo electrophysiology recording capabilities of MoRF probe implanted in CA1. **a**. An example wideband (0.1-6000Hz) extracellular trace showing theta oscillations (4-12Hz). **b**. An example wideband trace (0.1-6000Hz) capturing sharp wave-ripples (SPW-Rs, 100-250Hz). **c**. An example wideband trace (0.1-6000Hz) displaying multi-unit activity (MUA). **d**. Representative power spectral density (PSD) of smoothed local field potential (overlaid trace) across whole recording window in frequency bins (0-300Hz). **e**. Four isolated units. Each unit is color-coded to their respective autocorrelation patterns and spike waveforms. **f**. Representative 32-channel wideband extracellular recordings (0.1-6000Hz) from hippocampal CA1.

Additional electrophysiology recording was also performed with a 32-channel configuration (16 electrodes in the outer probe and 16 electrodes in the inner probe) in awake mice. Representative raw traces of all 32 channels recorded from CA1 are shown in Fig.3f.

### In vivo optogenetic stimulation

To validate the MoRF probe’s optogenetic stimulation capabilities, a 16-channel outer probe with a silica waveguide as the inner probe was implanted into awake mice with ChR2 expression restricted to CA1 pyramidal neurons. In these experiments, the probe was lowered slowly until detected electrophysiology signals exhibited oscillations verifying successful targeting of the CA1 layer. 473 nm laser pulses were delivered from a laser source modulated to output three short pulses with varying optical intensities. Fig.4a shows example electrophysiology traces in response to the three different light intensities. The lowest intensity optical pulse elicited no significant change in spike activity. The middle and high intensity optical pulses triggered more spikes. Note that the bottom raster plot demonstrates increasing neuron firing with higher optical stimulation intensity. Optogenetic stimulation of CA1 pyramidal cells elicited local high-frequency oscillations mimicking the native ripple as demonstrated previously in CA1^33^. Fig.4b indicates evoked high-frequency oscillations at the highest pulse intensity. These findings demonstrate the probe’s ability to modulate neural activity by controlling the optical input power.

**Fig. 4:**
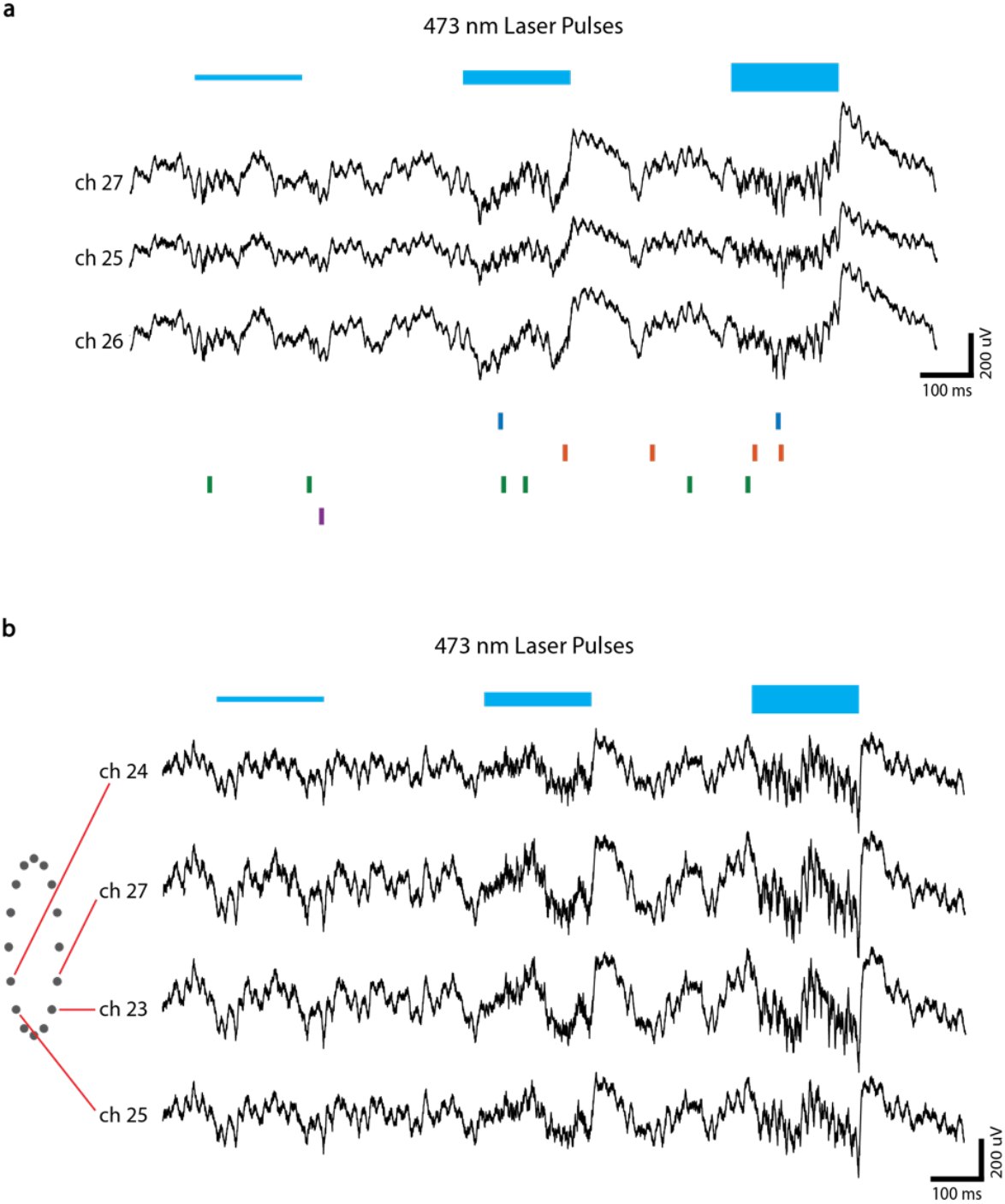
In vivo optogenetic stimulation capabilities of MoRF probe. **a**. Example traces of wideband (0.1-6000Hz) extracellular response to optical stimulation. **b**. Examples of optically evoked neural responses at various optical powers. No obvious response at the lowest optical stimulation intensity. Optically evoked neuron activity increased by varying the optical stimulation intensity, enabling induction of ripple-like high frequency oscillations at higher intensities. Oscillation amplitudes in channels 27 and 23 exceeded the other two channels, correlating with probe electrode sites as illustrated in the adjacent schematic.

## Discussion

Using thermal tapering, we developed the MoRF probe, a multifunctional polymer neural probe capable of switching functionalities and the configurations of its sensing element geometries. We demonstrated multiple possible configurations of the same device and validated its electrical and optical capabilities *in vivo*. We implanted the probe in awake mice to validate its performance and successfully recorded both 16 channel and 32 channel electrophysiology recordings, demonstrating its ability to record both LFP and single unit activities. We also conducted optical stimulation of opsin expressing neurons in the awake mouse, validating our device’s ability to evoke different neural activities with different optical stimulation intensities.

Existing neural probes’ sensing element densities, types, and configurations cannot be modified after probe fabrication, preventing scientists from tailoring devices for their specific experiments to achieve optimal performance. Here, for the first time, we designed and fabricated a general use neural interface platform that can be reconfigured by the researchers themselves. We accomplished this by designing an outer probe with a hollow channel in the center that allows the insertion and positional adjustments of inner probes with different functionalities. Such a design allows for considerable modularity, with its function configurations fully adjustable for optimal performance for each unique research scenario. Optical stimulation functions can be added by inserting a silica waveguide (easily obtainable from fiber optic suppliers) into the central channel, while drug delivery capabilities can also be incorporated by inserting a microfluidic tube in a similar fashion. Enhancing electrophysiology recording capabilities for situations requiring denser electrodes can also be accomplished by inserting an inner probe with multi-electrodes in the same manner, further multifunctionalities can also be incorporated by using a multifunctional inner probe. Additionally, our probe platform also allows researchers to configure the exact position of the sensing elements with respect to each other. By utilizing a microdrive to adjust the height difference between the inner and outer probes, researchers can control the distance between the outer and inner probe’s sensing elements. Such a feature enables straightforward multisite recording and stimulation in different target brain regions of interest. Although multisite stimulation and recording can also be realized by utilizing conventional probes with large recording areas or multiple probes inserted in different heights, such methods often have drawbacks such as increased cost and complexity, or decreased electrode density in areas of interest.

The MoRF probe’s functionalities are also not restricted to the ones demonstrated here. Its modular design means that any inner probe that can fit into the hollow channel of the outer probe is compatible. This feature allows it to be configured for a plethora of additional modalities such as chemical sensing, temperature sensing, and photometry. Additionally, the probe’s configuration and functions also have the potential to be adjusted post probe implantation, enhancing its chronic performance as well as offering scientists the ability to adjust device functionality to suit different phases of the experiment.

## Notes

### Competing Interest Statement

The authors have declared no competing interest.

### Funding

- US NIH: R01NS123069

